# Repurposing disulfiram (Tetraethylthiuram Disulfide) as a potential drug candidate against *Borrelia burgdorferi in vitro and in vivo*

**DOI:** 10.1101/842286

**Authors:** Hari-Hara SK Potula, Jahanbanoo Shahryari, Mohammed Inayathullah, Andrey Victorovich Malkovskiy, Kwang-Min Kim, Jayakumar Rajadas

## Abstract

Lyme disease caused by the *Borrelia burgdorferi* (*Bb or B. burgdorferi*) is a most common vector-borne, multi-systemic disease in USA. Although, most Lyme disease patients can be cured with a course of antibiotic treatment, a significant percent of patient population fail to be disease-free post-treatment, necessitating the development of more effective therapeutics. We previously found several drugs including disulfiram having with good activity against *B. burgdorferi*. In current study, we evaluated the potential of repurposing the FDA approved disulfiram drug for its *borrelia*cidal activity. Our *in vitro* results indicate disulfiram shows excellent *borrelia*cidal activity against both the log and stationary phase *B. burgdorferi*. Subsequent mice studies have determined that the disulfiram eliminated *B. burgdorferi* completely from hearts and urinary bladder by day 28 post infection, demonstrating the practical application and efficacy of disulfiram against *B. burgdorferi in vivo*. Moreover, disulfiram treated mice showed reduced expression of inflammatory markers and protected against histopathology and organ damage. Furthermore, disulfiram treated mice showed significantly lower amounts of total antibody titers (IgM and IgG) at day 21 and total IgG2b at day 28 post infection. Mechanistically, cellular analysis of lymph nodes revealed a decrease in percentage of CD19+ B cells and increase in total percentage of CD3+ T cells, CD3+ CD4+ T helpers, and naïve and effector memory cells in disulfiram-treated mice. Together, we demonstrate that disulfiram has the potential and could be repurposed as an effective antibiotic for treating Lyme disease in near future.

## Introduction

Lyme disease, a Zoonosis, is the most common reportable vector-borne disease in the United States and affects ∼300,000 persons annually in North America^1^, spread by the spirochete *Borrelia burgdorferi* sensu stricto (hereafter termed *B. burgdorferi or Bb*). The clinical manifestations of Lyme disease includes three phases^2^. Early infection consists of localized erythema migrans, followed within days or weeks by dissemination to the nervous system, heart, or joints in particular and subsequently resulting in persistent infections. Without antibiotics treatment, 60% of patients with Lyme disease in the United States develop arthritis, which may recur at intervals and last for months or years. Fewer patients (4 to 10%) suffer carditis, which is generally an early and nonrecurring manifestation of infection^3^. Antibiotic treatment usually with oral doxycycline, at the early localized stage of Lyme disease cures the disease in most patients^4^. However, 10 to 20 % of patients continue to experience major lingering symptoms, such as fatigue, musculoskeletal pain, and cognitive complaints, a condition known as post treatment Lyme disease syndrome (PTLDS)^5^. Several studies indicate that disseminated infection is not eradicated by conventional antibiotics such as tetracycline, doxycycline, amoxicillin or ceftriaxone in animal models tested like mice^6,7,8^, dogs^9^, ponies^10^ and in non-human primates.^11,12^ Several reports also showed that several antibiotics daptomycin and cefoperazone in combination with doxycycline or amoxicillin effectively eliminated *B. burgdorferi* persisters^13,14^. However, these antibiotic combination failed to act against *B. burgdorferi* biofilm forms^14^. Despite these findings, the PTLDS mechanisms are unclear and the plausible explanations for symptoms in animal models could be *B. burgdorferi* adapting multiple immune evasion mechanisms like alteration of highly immunogenic surface antigens^15,16^, inhibition of complement-mediated bacterial lysis^17,18^ that may render antibody response ineffective, there by supporting ongoing PTLDS. Therefore, based on these observations new mechanistic classes of antibiotics need to be developed to treat infections raising from these resistant forms of *B. burgdorferi*.

One approach to expedite the development of new antibiotics is to repurpose preexisting drugs that have been approved for the treatment of other medical conditions. Previously, we screened drugs (80% of them FDA approved, with a total of 4366 chemical compounds from four different libraries) with high activity against the log and stationary phase of *B. burgdorferi* by BacTiter-Glo™ Assay. Among them, disulfiram (Antabuse™), an oral prescription drug for the treatment of alcohol abuse since 1949, was found to have the highest anti-persister activity against *B. burgdorferi* ^19^. In addition, disulfiram and its metabolites are potent inhibitors of mitochondrial and cytosolic aldehyde dehydrogenases (ALDH)^20^. Recent U.S. clinical trials using repurposed disulfiram treatments include: methamphetamine dependence (NCT00731133); cocaine addiction (NCT00395850); melanoma (NCT00256230); muscle atrophy in pancreatic cancer (NCT02671890); HIV infection (NCT01286259) modulator of amyloid precursor protein processing (NCT03212599) and also recently initiated for previously treated Lyme disease (NCT03891667)^21^. In the area of infectious disease, disulfiram has been shown have antibacterial^22,23^, and anti-parasitic^24^ properties. Recently in a clinical setting, disulfiram appears to have conferred benefit in the treatment of a limited number of patients with Lyme disease and babesiosis^25^. Disulfiram is an electrophile that readily forms disulfides with thiol-bearing substances. *B. burgdorferi* possess a diverse range of intracellular cofactors (e.g., coenzyme A reductase)^26^, metabolites (e.g., glutathione), and enzymes (e.g., thioredoxin)^27^ containing thiophilic residues that disulfiram can potentially modify by thiol-disulfide exchange to evoke antimicrobial effects. Therefore, disulfiram has potential to inhibit *B. burgdorferi* metabolism by forming mixed disulfides with metal ions^28^ and it has been shown by our group previously that *B. burgdorferi* require zinc and manganese as co-factors for key biological processes^29^.

In the present study, we evaluated the antibacterial activities of disulfiram against log and stationary phases of *B. burgdorferi* in more detail. Furthermore, bactericidal activity of disulfiram *in vivo* was determined using the C3H/HeN mouse model of Lyme disease at early onset of chronic infection i.e. day 14 and day 21 post *B. burgdorferi* infection.

## Results

### Disulfiram as a potential antibiotic on log and stationary forms of *B. burgdorferi* B31

While our initial screen of disulfiram from four drug libraries is based on Bac titer-Glo assay^19^, we performed our preliminary disulfiram study of varying concentrations ranging from (100 µM to 0.625 µM) by Bac titer-Glo assay (Supplementary Figure 1A and 2A), which can only predict the cell viability based on quantitation of ATP present, but it cannot discriminate the inhibitory or bactericidal effects of the disulfiram. Further, we have performed the MICs/MBCs by gold standard micro dilution assay using the morphological evaluation methods like dark field direct cell counting and SYBR Green I/PI (Live/Dead) fluorescence microscopy counting in 48-well plate format (Figures 1 and 2). Here, in this study we evaluated disulfiram (dissolved in both DMSO and 30% hydroxypropyl β-cyclodextrin (from now on cyclodextrin) and doxycycline *in-vitro* sensitivity of spirochete and round body morphological forms, of *B. burgdorferi* B31 incubated for 4-5 days with different concentrations of drugs. Appropriate concentrations of cyclodextrin, DMSO and ultra-pure water in BSK medium were used as negative controls.

**Figure 1:**
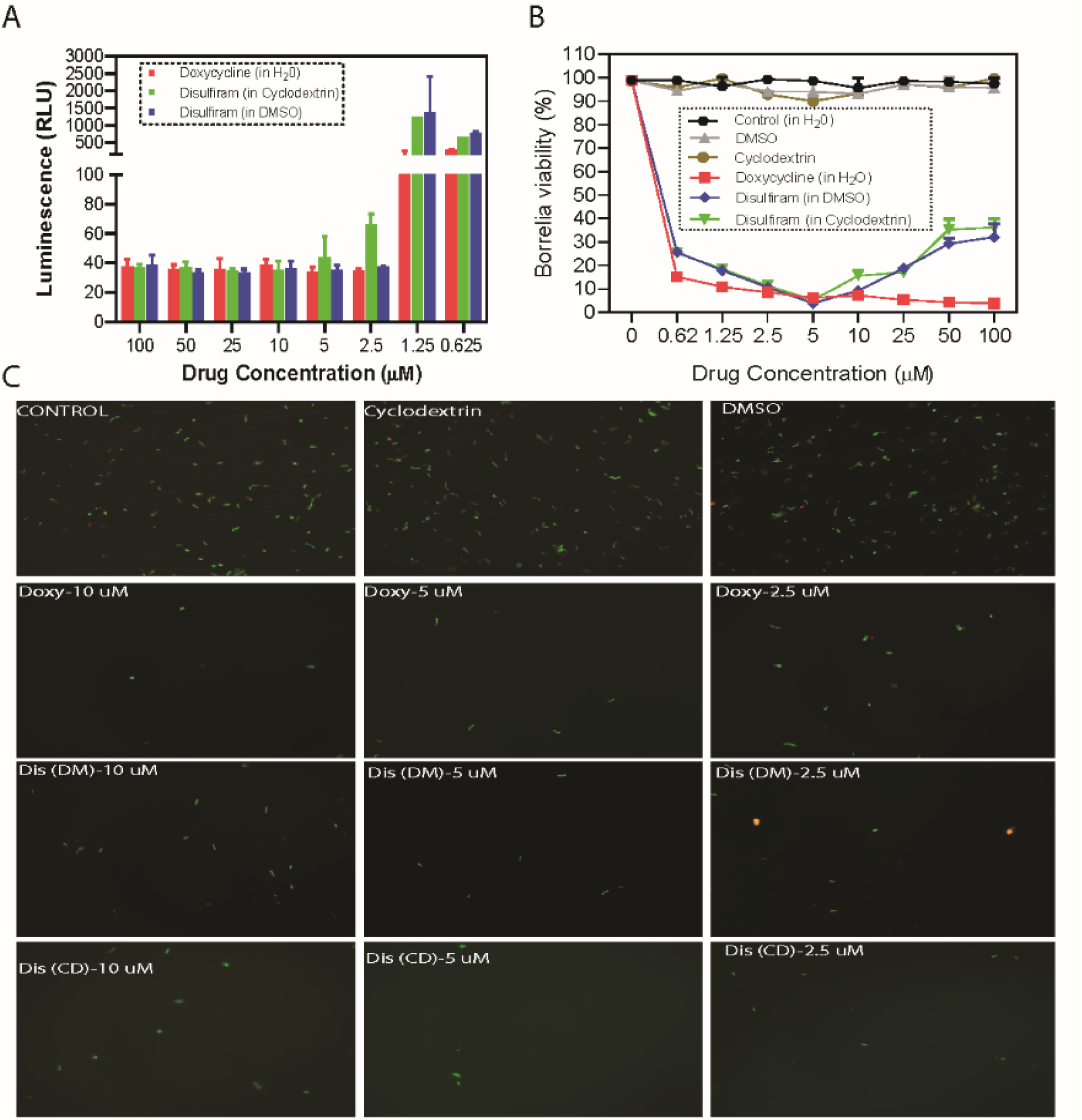
Evaluation of borreliacidal activity of disulfiram (in DMSO and cyclodextrin) with doxycycline as control. A 4 day-old, *B. burgdorferi* log phase culture of *B. burgdorferi* was incubated for four days with disulfiram (Dis-DM), disulfiram (Dis-CD) and doxycycline (Doxy) at the same drug concentrations of 100μM to 0.625μM respectively. After a five-day incubation, bacteria cell viability was assessed by **A**, Bac-titer glow assay. **B**, by direct counting using dark field microscopy and **C**, by SYBR Green-I/PI assay using fluorescent microscopy. Representative images were taken using SYBR green-fluorescent stain (live organisms) and propidium iodide red-fluorescent stain (dead organisms) at 20X magnification. All these experiments were repeated atleast three times. Error bars represent standard errors.

**Figure 2:**
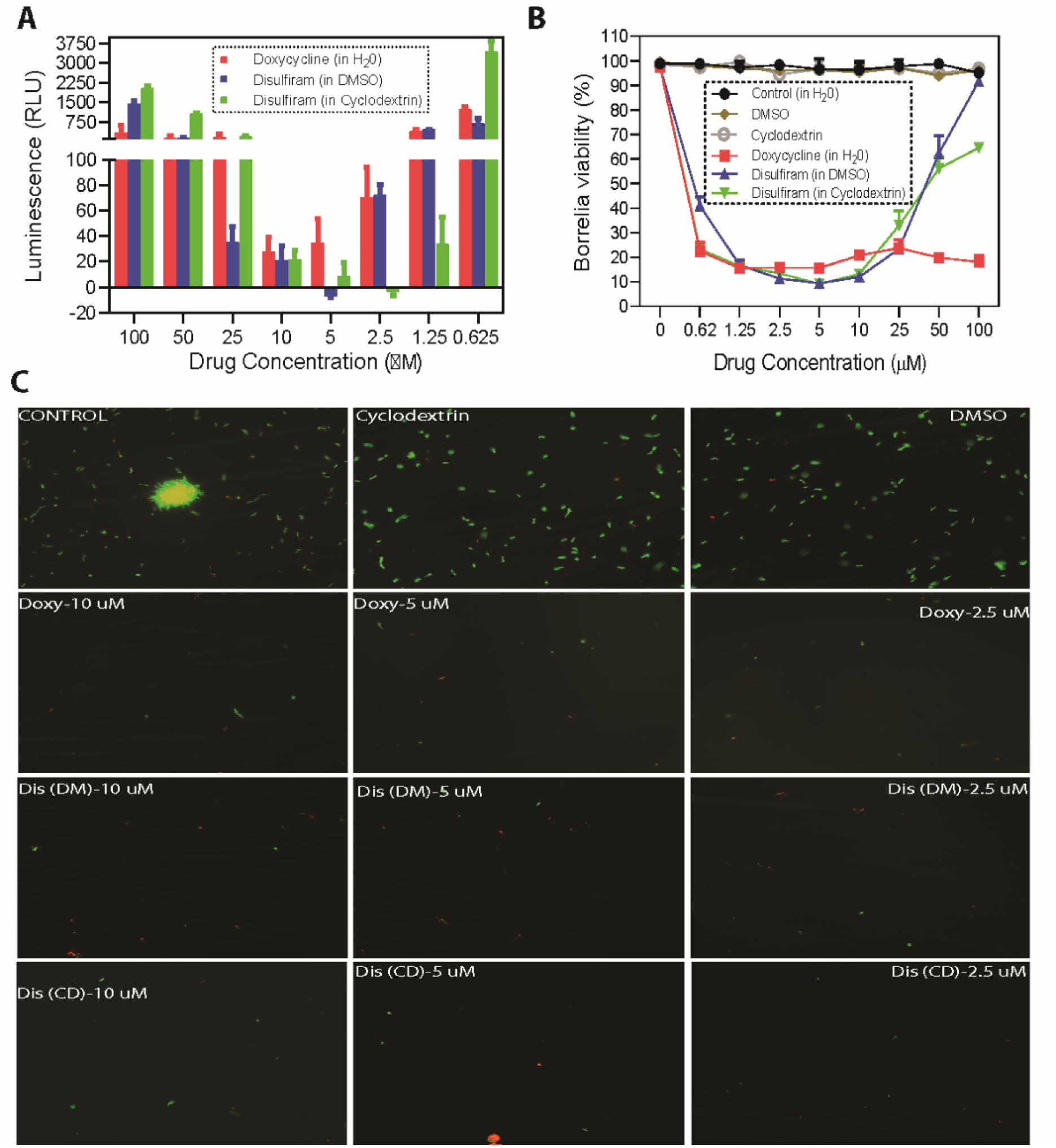
Evaluation of borreliacidal activity of disulfiram (in DMSO and cyclodextrin) with doxycycline as control. An eight day-old, *B. burgdorferi* log phase culture of *B. burgdorferi* was incubated for four days with disulfiram (Dis-DM) disulfiram, (Dis-CD) and doxycycline (Doxy) at the same drug concentrations of 100μM to 0.625μM respectively. After a five-day incubation, bacteria cell viability was assessed by **A**, Bac-titer glow assay. **B**, by direct counting using dark field microscopy and **C**, by SYBR Green-I/PI assay using fluorescent microscopy. Representative images were taken using SYBR green-fluorescent stain (live organisms) and propidium iodide red-fluorescent stain (dead organisms) at 20X magnification. All these experiments were repeated atleast three times. Error bars represent standard errors.

Our *in vitro* studies indicate that treatment with different concentrations ranging from 20 µM, 10 µM, 5 µM, 2.5 µM, 1.25 µM and 0.625 µM of disulfiram in DMSO and disulfiram in cyclodextrin drugs significantly eliminated log phase spirochetes ∼ 80-94% and stationary phase persisters ∼ 80-92% compared to the controls (Figures 1 and 2). More specifically, treatment with 5 µM (1.48 µg/ml) concentration of disulfiram in DMSO and disulfiram in cyclodextrin drugs significantly eliminated log phase spirochetes ∼ 94% and treatment with 5 µM (1.48 µg/ml) concentration of disulfiram in DMSO and disulfiram in cyclodextrin drugs significantly eliminated stationary phase spirochetes ∼ 92% and ∼ 90% respectively (Figure 2A and 2B). In our study also, as reported earlier^13^ doxycycline was significantly able to reduce the viability of log phase *B. burgdorferi* by ∼90-97% compared to the control (Fig. 1A and 1B). However, doxycycline treatment had no significant effect on the cells in the stationary phase cultures as observed by increasing proportion of viable cells after antibiotic exposure compared to the control (Figure 2B and 2C). However, at high concentration (ranging from 50 µM-100 µM) lose efficacy and shows reduced bactericidal activity with increase in concentration of disulfiram drug in DMSO or cyclodextrin. These results are specific since treatment with ultra-pure water or cyclodextrin or DMSO in BSK medium did not significantly reduce the viability of the spirochete rich log phase culture and persisters rich stationary cultures compared to the control observed in both direct dark field and SYBR Green-I /PI based fluorescent microscopy counting respectively (Fig. 1A and 2A).

To validate our preliminary results, *B. burgdorferi* B31 log and stationary forms were further evaluated *in vitro* for disulfiram in DMSO and disulfiram in cyclodextrin drugs sensitivity by a fluorescent microscopy counting using SYBR Green-I (live cells stain green) and Propidium Iodide (dead cells stain red). Consistently, the 5µm (1.48 µg/ml) concentration of disulfiram in DMSO and disulfiram in cyclodextrin drugs significantly reduced log phase spirochetes by ∼ 94 %, but in the remaining 6 % of the population, ∼ 4 % were stained green for live while ∼ 2 % were stained red for dead (Figure 2B). While doxycycline significantly reduced the log phase viability by ∼ 97%, it did not reduce the stationary phase viability of *B. burgdorferi* (Figure 2B). However, most interestingly treatment with 5 µm (1.48 µg/ml) concentration of disulfiram in DMSO and disulfiram in cyclodextrin drugs significantly eliminated stationary phase spirochetes ∼ 92 % and ∼ 90 % respectively but in the remaining 8 % and 10 %, ∼ 6 % and 8 % were live while ∼ 2 % were dead (Figure 2B). These results agreed with the dark field microscopy counting.

At low concentrations (ranging from 10 µM-0.625 µM), the disulfiram in DMSO and disulfiram in cyclodextrin drugs concentration response profile is sigmoid. In contrast, at higher concentrations (ranging from 25 µM-100 µM), the disulfiram drugs lose efficacy, exhibiting the U-shaped or bell curve observed in the Figures 1 and 2. We attribute this loss in activity to the drugs being in colloidal form. This was further analyzed by the Dynamic light scattering (DLS) and Atomic force microscopy based techniques.

### Disulfiram forming aggregates at high concentration was shown by DLS study and AFM based imaging

Dynamic light scattering (DLS) technique was used to study the aggregation of disulfiram. Supplemental figure 1 indicates a variation in the average count rate of the particles with increasing concentrations of disulfiram prepared from DMSO and CD stock solutions. The samples from DMSO preparation showed a linear increase in the average count rate with respect to the concentration. However, the slope of the increase changed above 10 µM indicating a critical aggregation concentration (CAC). The results from the CD preparation showed a non-linear trend with a break at 10 µM consistent with the CAC results from DMSO preparation.

Atomic force microscopy-based techniques helped us to further evaluate the small volume (10 µL) of liquid sample aliquots and fast drying of the highly spread droplets on hydrophilic substrates allow to assess the real disulfiram particle dimensions with minimal contribution of secondary sample aggregation due to local increase in its concentration due to drying. In supplemental figure 2 observed that very few aggregates were formed for DMSO samples for all concentrations (supplemental figure 2; E-H). The smaller particles are less than 1 nm in height.

For cyclodextran particles, we can see larger particles that are quite wide, but also quite flat – only 10 nm high, on average. These are likely particles formed due to sample drying. However, for smaller particles, a crossover can be seen from 25 to 10 µM (supplemental figure 2; A-D). In the former sample, we can still observe them (white arrows), but not in the latter, which is only 2.5 times less concentrated. Thus, this can be proof of sample aggregation, which starts to be substantial only above 20 µM, as evident from our DLS data.

### Disulfiram treatment reduces the *B. burgdorferi* burden in tissues following dissemination in infected C3H/HeN mice

Following-up on the potent killing *in vitro* activity of disulfiram against *B. burgdorferi*, we examined its efficacy *in vivo* immunocompetent C3H/HeN^30^ mice and compared with doxycycline. To better compare the efficacy, disulfiram (75 mg/kg of bodyweight) was introduced intra peritoneally on day 14 and day 21 (to consistently develop persistent infection and carditis^31^) in to post infected C3H/HeN mice at 75 mg/kg of body weight every day for 5 days (Figure 3A). Mice were sacrificed after 48 hours of last dose to collect tissue samples at both time points (day 21 and day 28 post drug treatment). A whole ear and heart base were used for cultivation and analyzing the pathogen loads. Spleen, bladder, ear and heart were collected for whole-DNA extraction and Q-PCR analysis (Table 1 and Figure 3A). The *flab* gene PCR positivity represent either live *Bb* or components that had not been cleared from tissues. In the day 14 post disulfiram treatment group, 3 out of 5 mice were *flaB* gene PCR positive from heart, bladder and ear tissues (Table1 and Figure 3B). Whereas in the doxycycline treatment (50 mg/kg) group, 3 out of 5 mice were positive by PCR from heart and ear tissues but none positive for PCR from bladder tissues (Figure 3B). Similar results observed in cultured ear and heart tissues of respective groups (data not shown). The disulfiram treatment group had statistically significant lower number of *B. burgdorferi* compared to untreated infected controls (Table 2 and Figure 3B). On the other hand, in the day 21 post disulfiram treatment group, 2 out of 4 mice were *flaB* gene PCR positive from ear and rest of the tissues heart and bladder were PCR negative (Figure 3B). Whereas in the doxycycline treatment group, 4 out of 4 mice ears were PCR positive and 1 out of 4 mice bladder was PCR positive but none positive for PCR from heart (Table 2 and Figure 3B). Similar results observed in cultured ear and heart tissues of respective groups (data not shown). These results showed an overall better efficacy *in vivo* C3H/HeN mouse to restrict the further growth and dissemination of *B. burgdorferi*.

**Table 1:**
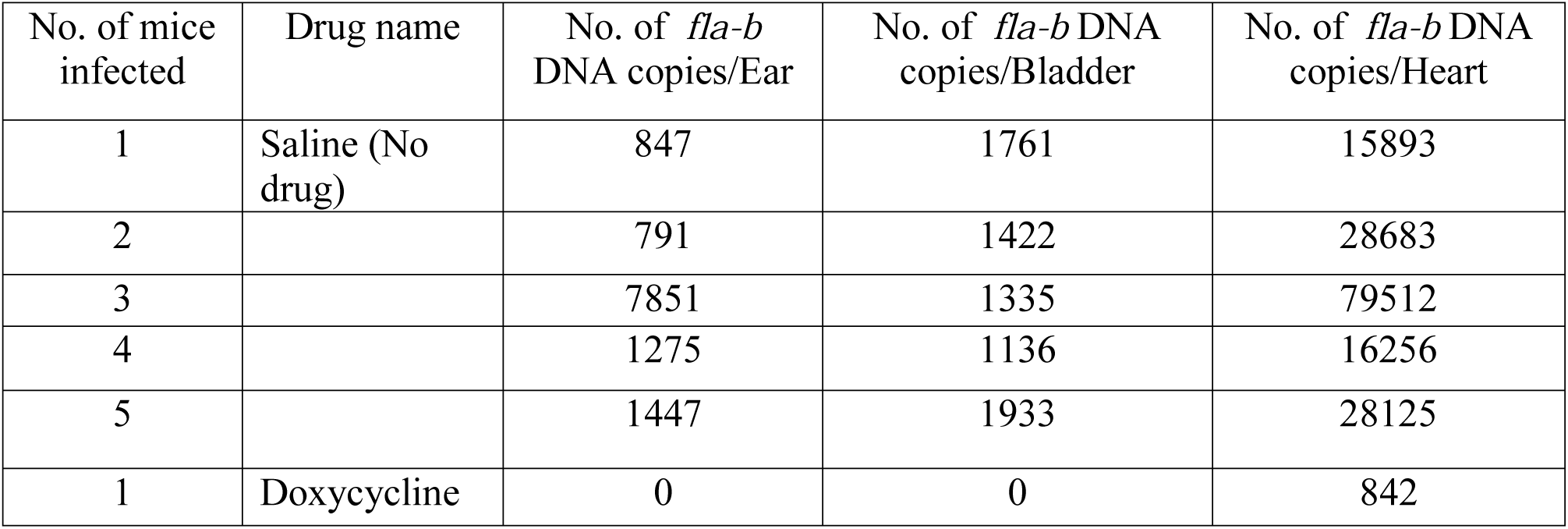
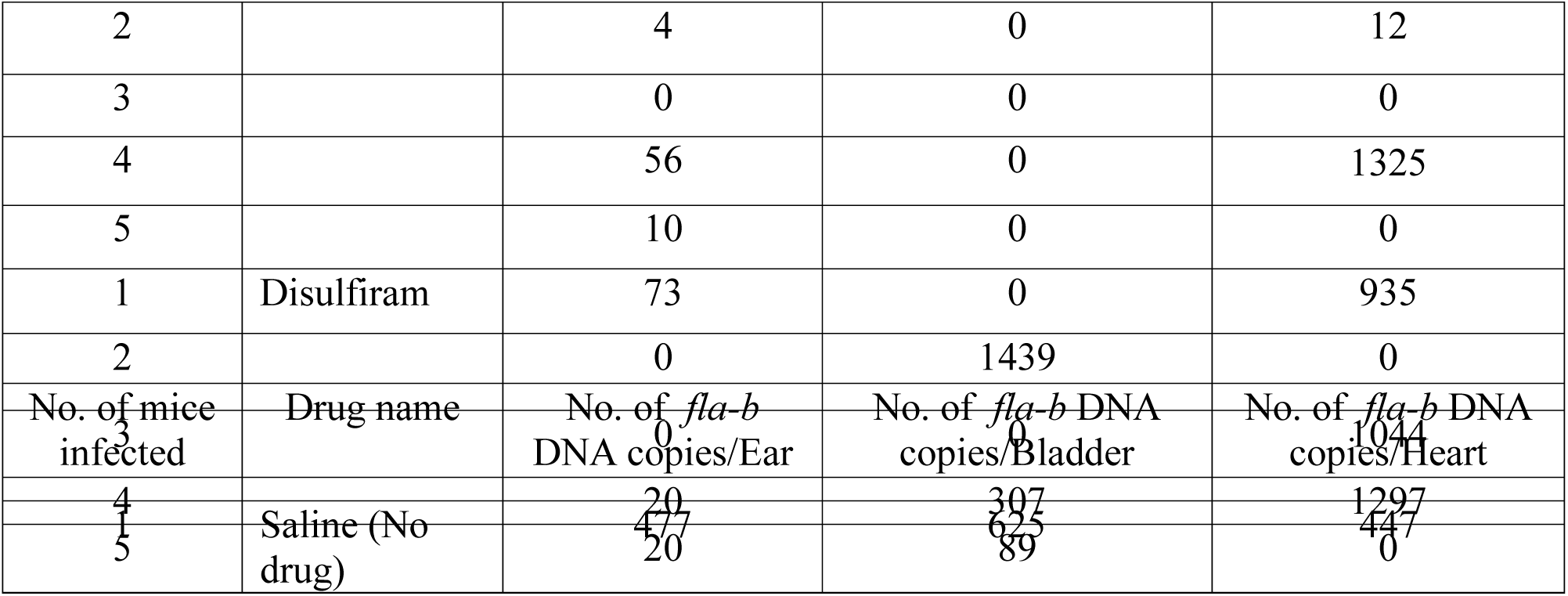
*In vivo* efficacy of drugs against *B. burgdorferi* in C3H/HeN mice. After 14 days of *B.burgdorferi* infection, C3H/HeN mice were treated with following drugs once per day for 5 days (Doxycycline – 50 mg/kg and Disulfiram – 75 mg/kg). The whole DNA was extracted from urinary bladder, ear and heart and further analyzed data by qPCR.

**Table 2:**
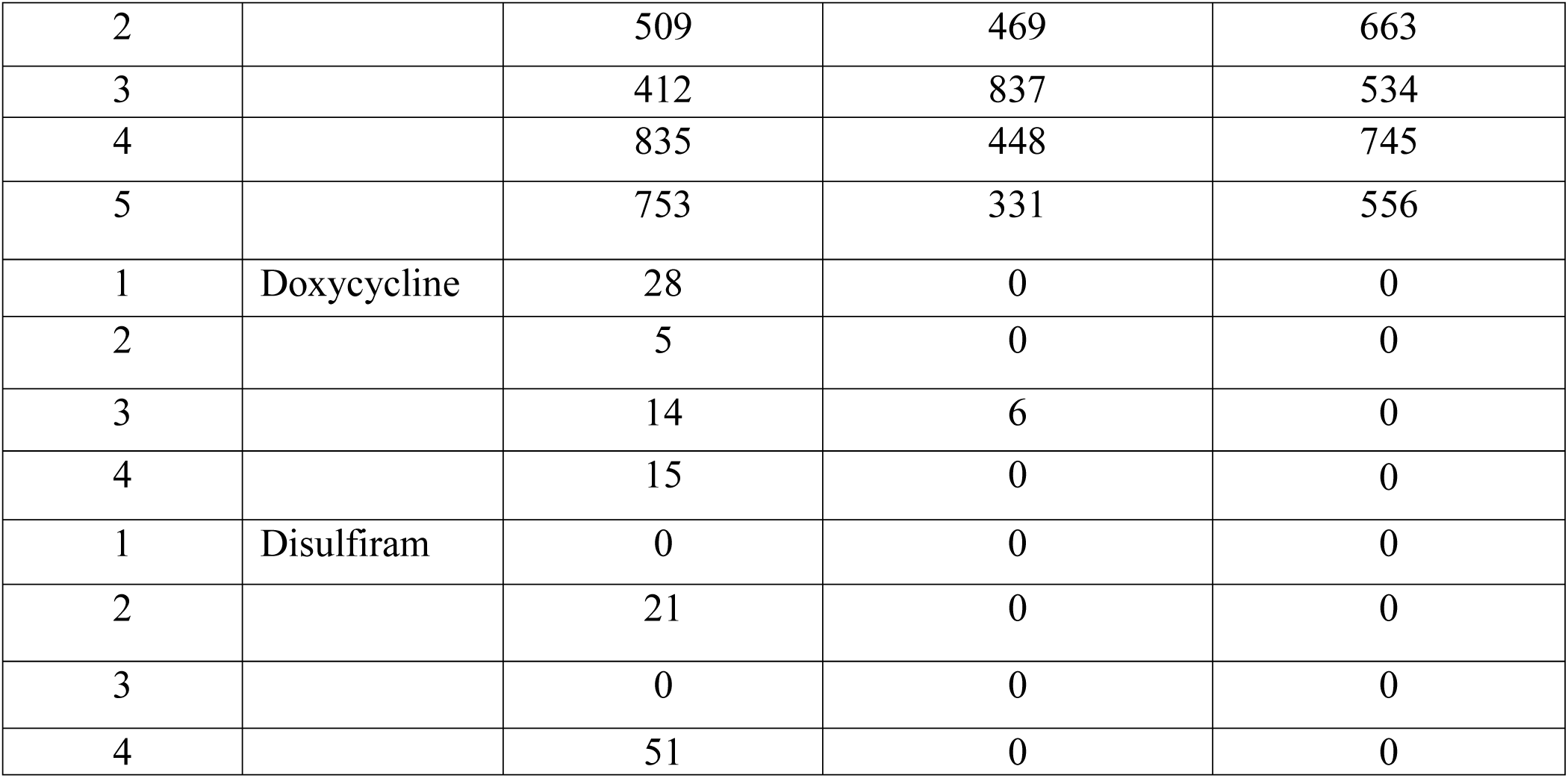
*In vivo* efficacy of drugs against *B. burgdorferi* in C3H/HeN mice. After 21 days of *B.burgdorferi* infection, C3H/HeN mice were treated with following drugs once per day for 5 days (Doxycycline – 50 mg/kg and Disulfiram – 75 mg/kg). The whole DNA was extracted from urinary bladder, ear and heart and further analyzed data by qPCR.

|  |  |  |  |  |
| --- | --- | --- | --- | --- |
| 2 |  | 509 | 469 | 663 |
| 3 |  | 412 | 837 | 534 |
| 4 |  | 835 | 448 | 745 |
| 5 |  | 753 | 331 | 556 |
| 1 | Doxycycline | 28 | 0 | 0 |
| 2 |  | 5 | 0 | 0 |
| 3 |  | 14 | 6 | 0 |
| 4 |  | 15 | 0 | 0 |
| 1 | Disulfiram | 0 | 0 | 0 |
| 2 |  | 21 | 0 | 0 |
| 3 |  | 0 | 0 | 0 |
| 4 |  | 51 | 0 | 0 |

**Figure 3:**
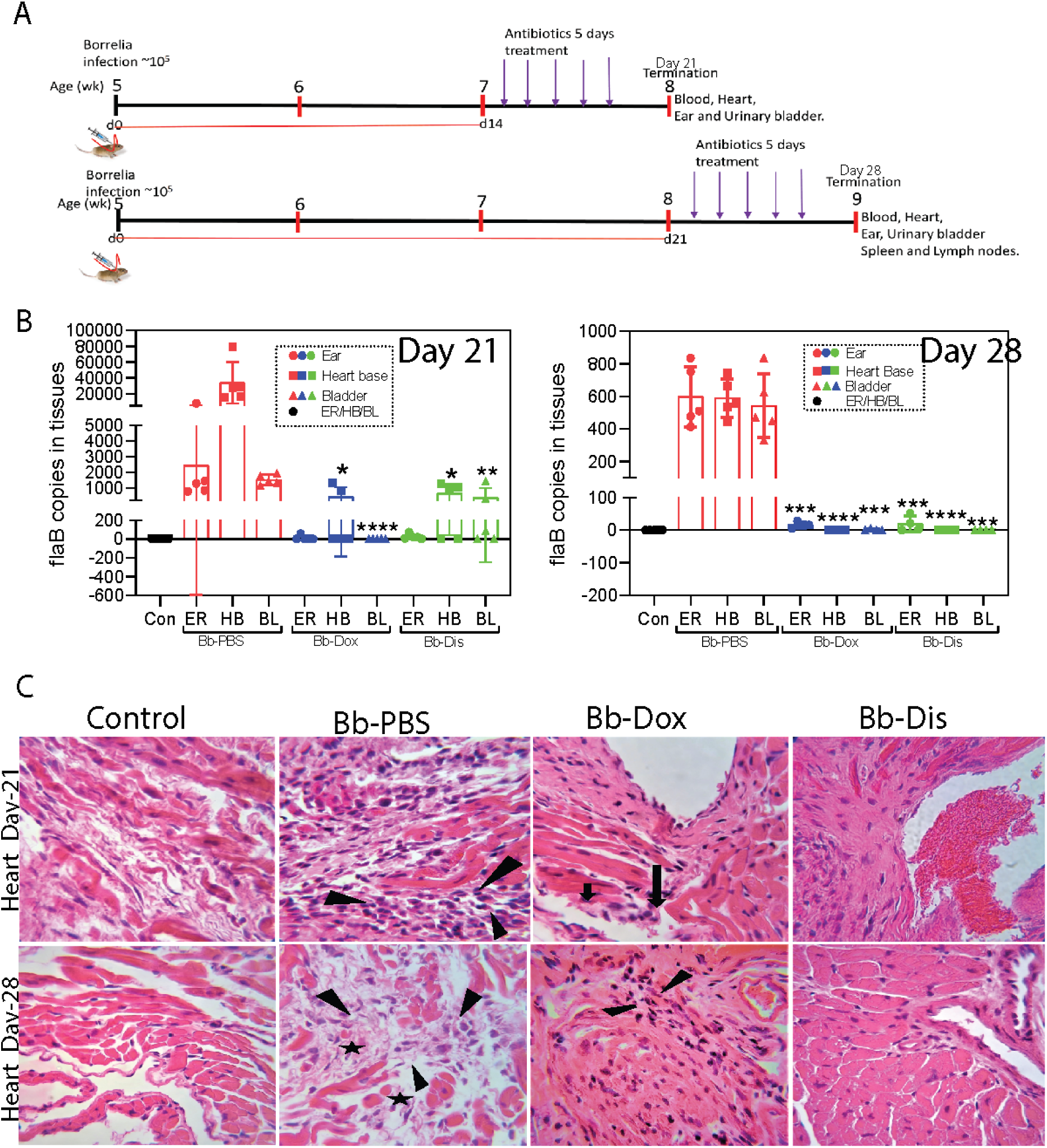
Borrelia loads in various tissues after C3H-HeN mice infection followed by disulfiram or doxycycline antibiotics treatment. **A.** Antibiotic treatment/borrelia infection schedule: groups of 5 week old C3H mice (n = 30) were infected subcutaneously above the shoulders with mid log phase 10^5^ *B. burgdorferi.* Infected groups were received intraperitoneal antibiotics [doxycycline (n=5), disulfiram (n=5) and PBS (n=5)] in two different time points; day-14 post infection and day-21 post infection. Uninfected groups of mice were kept as controls (n=10). **B.** Necropsy at the end of day-21 and day-28, ears, hearts and urinary bladders were collected for determination of the number of *Borrelia* flab per ul of sample by qPCR. **C.** Photomicrographs (40X) of hematoxylin and eosin stained heart sections; arrows depict the mono nuclear leucocyte infiltrates. Statistics by unpaired t test with Welch’s correction between drug treated group versus infected group. *p < 0.05, ** p < 0.01, *** p < 0.001, ****p <0.0001.

### Disulfiram treatment decreases disease pathology and further reduces inflammatory markers in the heart of *B. burgdorferi* infected C3H/HeN mice

In Lyme borreliosis, heavy inflammatory infiltrates dominated by mono or polymorphonuclear leukocytes are typically found at lesion sites^32^. We performed histopathology analysis of heart in both the day 14 and day 21 post disulfiram treatment group mice and found normal features of aorta, valves and few to no mononuclear leukocytes inflammation in the myocardium which signifies inactive carditis than the infected mice (Figure 3C and supplemental figure 3), whereas the doxycycline treatment group mice specifically in day 21 showed mild to moderate level of mononuclear leukocytes inflammation in the aorta and valves which signifies active carditis (Figure 3C and supplemental figure 3) than the infected untreated mice which showed transmural infiltration of mononuclear leukocytes in the aorta and valves signifies severe active carditis (Figure 3C and supplemental figure 3).

In infectious diseases, a hallmark of inflammatory tissue reactions is the recruitment and activation of leukocytes. Chemo and cytokines play a pivotal role in mediating these events. We further determined whether disulfiram treatment alters the inflammatory responses in the heart at both day 14 and day 21 post infection, we evaluated *B. burgdorferi* induced myocardial inflammation by quantification of mRNA transcription of CxCL1 (KC), CxCL2 (MIP-2), CCL5 (RANTES), TNFα, IFNγ, IL-10, IL-1β, iNOS and NOS-2 by qRT-PCR. In the day 14 post disulfiram treatment group, levels of MIP-2, TNFα, IFNγ and IL-10 were significantly lower (reached normal levels) relative to infected untreated mice (Figure 4). More specifically, IL-10 levels were reduced to 60-fold (Figure 4C), while MIP-2, TNFα, and IFNγ reduced to 10-fold (Figures 4 A, D&E). There was no change in NOS2 and iNOS levels (Figures 4 G&H). On the other hand, in the day 21 post disulfiram treatment group, levels of MIP-2, RANTES, TNFα, IFNγ, IL-1β, and IL-10 were significantly lower (touched to normal levels) relative to infected untreated mice (Figure 4), while iNOS and NOS-2 levels were significantly higher relative to infected untreated mice (Figure 4 G&H). More specifically, IL-10 levels were reduced to 100-fold (Figure 4 C), and other cytokines like MIP-2, TNFα, IL-1β, and IFNγ levels were reduced ten to sixty-fold (Figures 4 A, D, F&E). While NOS-2 and iNOS, which have role in immune regulation and tissues remodeling, were significantly higher compared to infected untreated mice (Figures 4 G & H). These results indicate that disulfiram affects the regulation and/or balance of Th1 (MIP-2, RANTES, TNFα, IFNγ and IL-1β), Th2 (IL-10) and protective Macrophage M1 (NOS2, iNOS) responses to *B. burgdorferi* at day 21 and day 28 post infection. However, doxycycline treatment group have only reduced few cytokines like IL-10, TNFα, and IFNγ at day 14 and 21 post treatment (Figures 4 C, D, & E). While NOS-2 levels elevated at day 21 and MIP-2 levels reduced at day 28 post infection (Figures 4 G & H).

**Figure 4:**
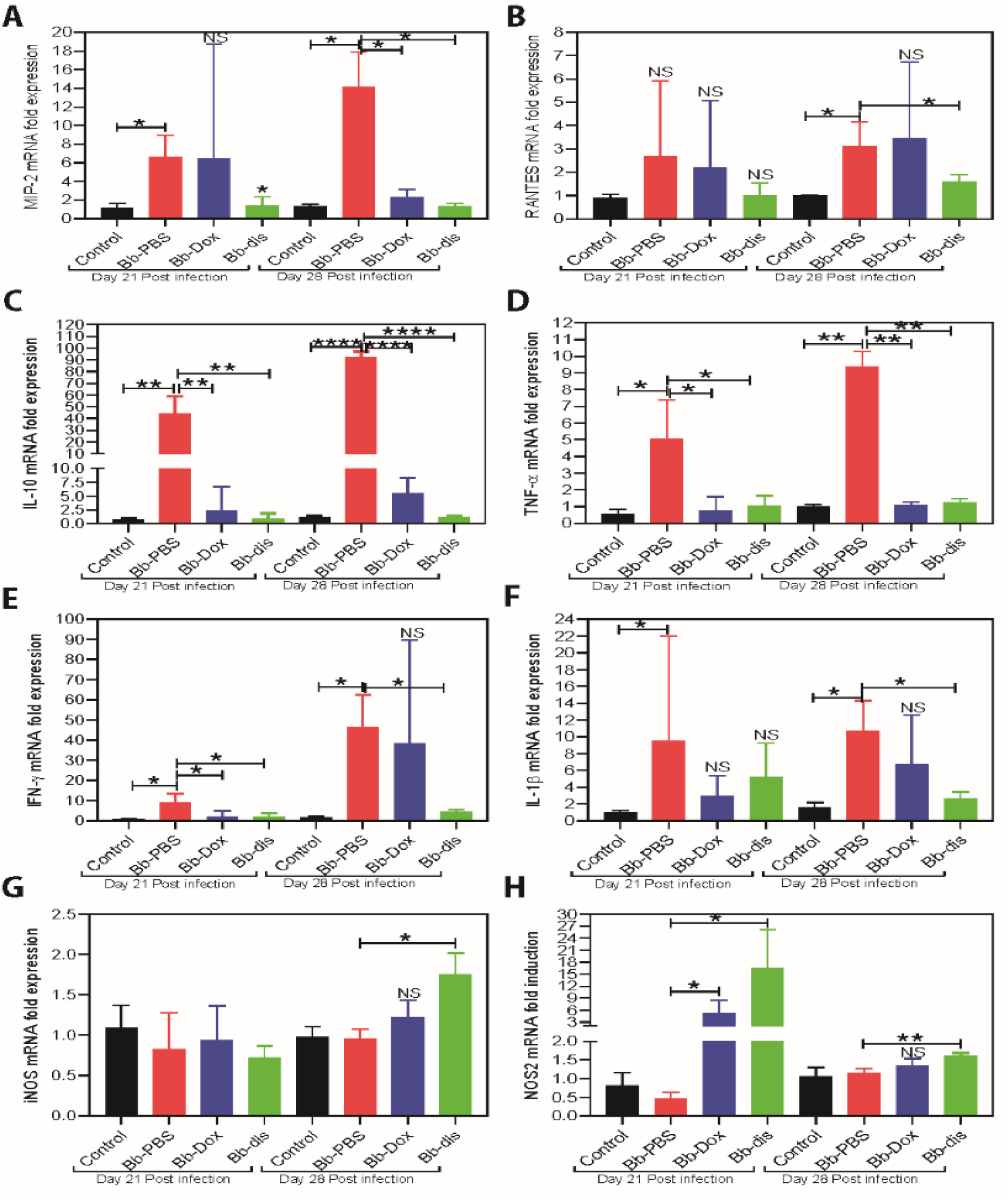
Measurement of immunomodulators in hearts of C3H/HeN mice with or without antibiotics treatment by RT-PCR. RT-PCR of pro-inflammatory transcripts (MIP-2, RANTES, TNF-α, IFN-γ and IL-1β) and important protective immunoregulatory transcripts (IL-10, iNOS and NOS-2) in heart. Statistics by unpaired t test with Welch’s correction between control versus infected and also between drug treated group versus infected group. *p < 0.05, ** p < 0.01, *** p < 0.001. NS means not significant.

### Disulfiram treatment reduces antibody titers in the *B. burgdorferi* infected mouse

We next sought to determine whether disulfiram treatment affects antibody development during day 14 and day 21 post *B. burgdorferi* infection, we measured the serum levels of each subtype of total immunoglobulins using an ELISA. The results at day 21, showed that the total amount of IgM and IgG levels were significantly lower in disulfiram treated mice compared infected control mice (Figure 5A). Among the IgG subtypes, total IgG1 levels were significantly lower than the infected control mice (Figure 5A). However, there was no effect on other IgG subtypes like IgG2a, IgG2b and IgG3 (Figure 5A). While at day 28, only trend towards lower IgG levels observed but were not statistically significant (Figure 5B). However, IgG2b levels were significantly lower in disulfiram treated mice and no effect on other IgG subtypes like IgG1, IgG2a and IgG3 (Figure 5B). Whereas doxycycline treatment group does not show any reduction of antibody titers at day 21 and day 28 respectively (Figures 5A & B). These data suggest that the disulfiram treatment might induced development of antibody subtypes very efficiently and affect IgG class switching, which may represent a contributing factor in lowering *B. burgdorferi* titers at day 21 and clearance of *Bb* more efficiently at day 28. However, we cannot exclude the fact that there is a possibility that B cells expressing different immunoglobulin isotypes are selectively expanded.

**Figure 5:**
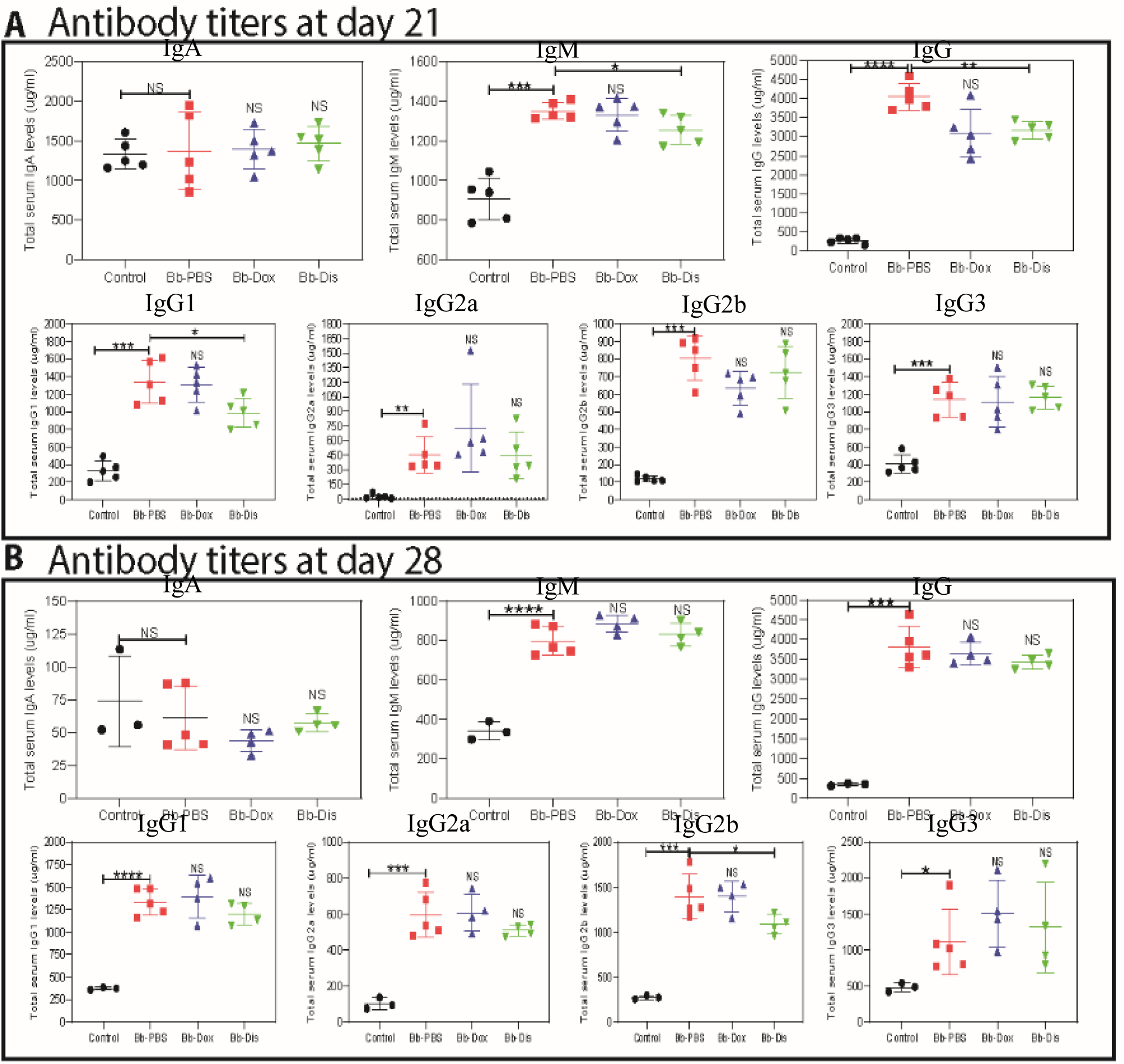
Antibody response in the serum with and without antibiotics treatment. A. Total concentration of IgA, IgM, IgG, IgG1, IgG2a, IgG2b and IgG3 antibodies day 21 post-infection and B. Total concentration of IgA, IgM, IgG, IgG1, IgG2a, IgG2b and IgG3 antibodies day 28 post-infection which were quantified by ELISA. Statistics: unpaired t test with Welch’s correction between controls versus infected and between drug treated group versus infected group. *p < 0.05, ** p < 0.01, *** p < 0.001. NS means not significant.

### Disulfiram reduces lymphoadenopathy in *B. burgdorferi* infected C3H/HeN mice

Lymphadenopathy, a hall mark of acute Lyme borreliosis^33^ manifestation is characterized by increased cellularity and the accumulation of large pleomorphic IgM- and IgG-positive plasma cells. To determine whether disulfiram treatment reduces the lymph node enlargement, at day 28 we collected peripheral (axillary, brachial, cervical and inguinal) lymphnodes (pLNs) and determine the cell number counts followed by analyzing B and T cell populations by flow cytometry. In disulfiram treatment mice, total lymphocytes of pLNs were statistically reduced in comparison to infected control mice (Figure 6). Doxycycline treatment mice also shown similar result. Further, our pLNs FACS analysis of disulfiram treatment mice had significant decrease of the percentages of CD19+ B cells, and significant increase of the percentages of CD3+ T cells in comparison to infected control mice (Figure 6). Further among the CD3+ subsets, CD3+ CD4+ helper T cells and CD3+ CD8+ cytotoxic T cells were not affected in comparison to infected control mice (Figure 6). However, when we compare naïve uninfected mice with all three infected groups (infected PBS treated, infected doxycycline treated and disulfiram treated) showed significant decrease of the percentages of CD3+ CD8+ cytotoxic T cells (Figure 6), and significant increase of the percentages of CD3+ CD4+ helper T cells (Figure 6). Another hallmark of effective and long-term protection is the generation of memory T cells. They provide an efficient immune response on pathogen re-exposure^34^. We further analyzed CD4^+^ T helper subsets by labeling naïve (CD62L^+^), early effector (CD62L^−^/CD44^−^), effector (CD44^+^) and memory T cells (CD62L^+^/CD44^+^). Analysis of helper T cells in comparison to naïve uninfected mice revealed that disulfiram treatment mice led to a significant increase of early effector/effector and memory T cells and to a significant decrease of naïve T cells in pLNs (Figure 6). Similar trend was observed in infected PBS treated and infected doxycycline treated mice (Figure 6).

**Figure 6:**
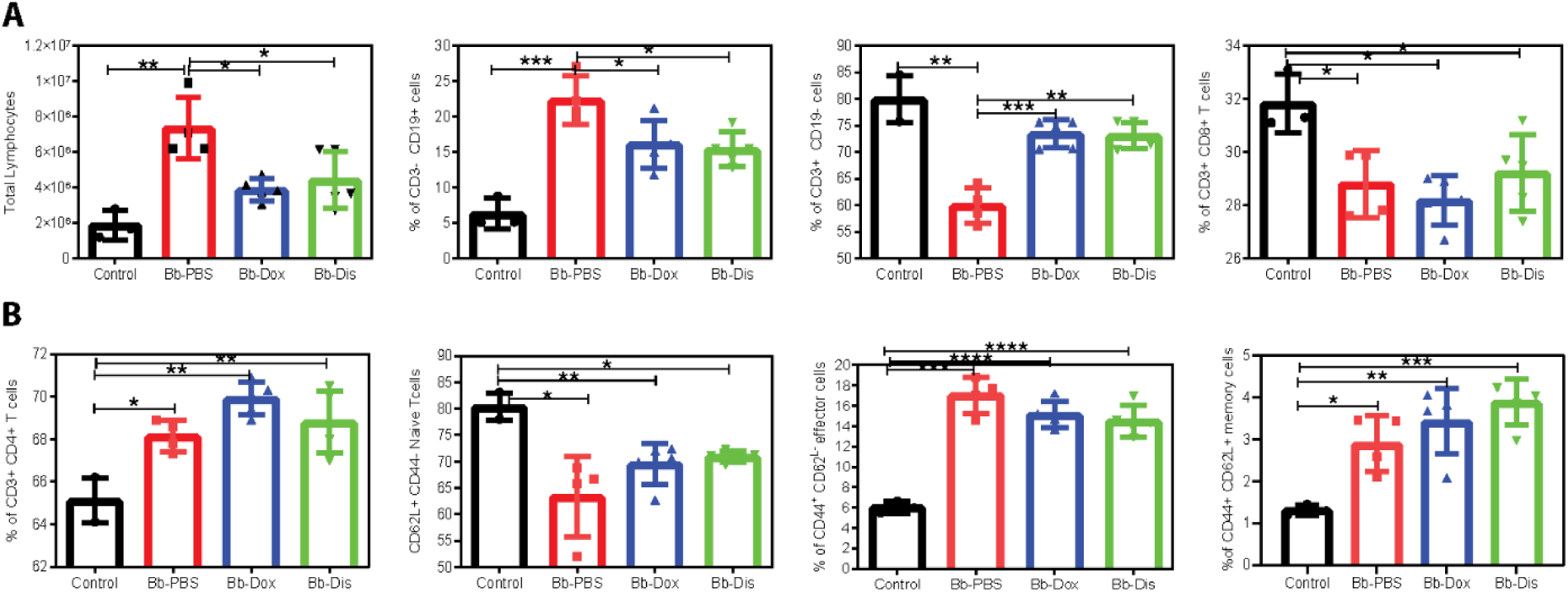
Percentage of B cells, T cells, CD8+ cytotoxic T cells, CD4+ helper T cells, naïve, effector and memory CD4+ T cells in lymph nodes. Flow cytometric analysis of immune cells isolated from peripheral lymph nodes from uninfected and infected mice treated with antibiotics. Cells were labelled with anti-CD19, anti-CD3, anti-CD4, anti-CD8, anti-CD44 and anti-CD62L lineage surface markers. Statistics unpaired t test with Welch’s correction between control versus infected and between drug treated group versus infected group. *p < 0.05, ** p < 0.01, *** p < 0.001. NS means not significant.

## Discussion

Since antibiotics are the top-of-the-line options to treat infections, there remains a dire need and a practical approach to bring more efficient antibiotics to clinic. The repurposing of FDA approved antibiotics through fast-track approvals can be an excellent solution. In this current study, we evaluated the borreliacidal potential of FDA approved drug disulfiram *in vitro* and *in vivo* based on our previous high-throughput screening hits^19,35^. We performed preliminary *in vitro* antimicrobial assays by Bac-titer glo assay with wide range of disulfiram concentrations (0.625 µM to 100 µM). Later, we confirmed the preliminary results by comparing antimicrobial effect of disulfiram to doxycycline and used most reliable quantitative methods performed to establish the bactericidal activity^36,37^. Disulfiram in both soluble forms (DMSO or cyclodextrin) inhibited the the growth of *B. burgdorferi* strain B31 at an MIC^90^ range of 0.74 to 2.97 µg/ml in case of log phase cultures (∼94%) and stationary phase cultures (∼90%) at 1.48 µg/ml, respectively (Figures 1 and 2), and with MBC varying from 1.48 µg/ml to 2.97□µg/ml for log and stationary phase cultures. The immediate deceleration in log and stationary phases of *B. burgdorferi* growth on low dose of disulfiram treatment is attributed to the rapid cleavage of disulfiram by thiophilic residues in intracellular cofactors like coenzyme A reductase^26^, enzymes like thioredoxin^27^, metal ions like zinc and manganese^29^, and cofactors of *B. burgdorferi*, which are hypothesized to instigate an abrupt halt in *B. burgdorferi* metabolism, thus evokes killing of *B. burgdorferi*. A similar mechanism of action is proposed for pathogens like Giardia, Bacillus, drug resistant Mycobacterium, and multidrug resistant Staphylococcus ^38,22,23,39,40^.

Disulfiram is an oral medication that is approved by the U.S. Food and Drug Administration (FDA) for administration of up to 500 mg daily^41^. Pharmacokinetic studies in humans has shown that disulfiram has a half-life (t_1/2_) of 7.3 h and a mean plasma concentration of 1.3 nM, although significant intersubjective variations are noted^42^. The toxicity of both disulfiram and its metabolites have also been broadly investigated in cell and animal studies, which yielded no evidence for teratogenic, mutagenic, or carcinogenic effects^43^. DMSO proved to be low dose toxic *in vivo*^44^ so, we have used non-toxic cyclodextrin^45,46^ as a solubilizing agent for disulfiram *in vivo* studies. Based on these observations, we conducted our preliminary *in vivo* mouse efficacy studies by administering (I.P) low dose of 10 mg/kg of body weight disulfiram to infected C3H/HeN mice for 5 days and found that these mice could not able to clear the *B. burgdorferi* from tissues (unpublished data). However as shown in the current study, when we repeated *in vivo* C3H/HeN mouse efficacy studies by administering (I.P) 75 mg/kg of body weight disulfiram to infected mice for 5 days, all infected mice either reduced or cleared the bacteria in most of the tissues at 21 and 28 post infection (Tables 1, 2 and Figure 3). C3H mice develop bradycardia and tachycardia beginning on day 7 through 60 days after *B. burgdorferi* inoculation and reaches severe inflammation particularly in C3H mice on day 15 to 21 post infection^31^. So, we have chosen C3H/HeN mouse model for our efficacy studies and day 14 or day 21 post infection as time points for antibiotics treatments. Lyme carditis, a macrophage-mediated pathology not directly influenced by *B. burgdorferi* specific antibodies, but by inflammatory micro environment created by mRNAs for proinflammatory Th1 cytokines (IL-1β, TNF-α, and IFN-γ), Th2 (IL-10), and other M1/M2 protective macrophage polarizing factors like iNOS and NOS2 derived from macrophages and T cells^47,48,49^. Similarly, chemokines like MIP-2 (macrophage inflammatory protein 2), KC, and RANTES (regulated upon activation, normal T cell expressed and secreted) preferentially attract monocytes and lymphocytes significantly contributing to the inflammation and tissue damage in Lyme disease^50^. We have shown that in disulfiram treatment mice there is a significant reduction in the infiltration of leucocytes in the heart wall and leads to no inflammation (inactive carditis) compared to doxycycline treated group (active mild carditis) and PBS infected group (active severe carditis) at day 21 or day 28 post infection (Figure 3). Which implies that disulfiram treatment reduced the inflammatory microenvironment by reducing the inflammatory chemokines (MIP-2 and RANTES), and cytokines (IL-10, IL-1β, TNF-α, and IFN-γ) and further reduces the disease severity in heart. Macrophage phenotype is flexible, and once the infection is cleared and a more anti-inflammatory environment is created, these inflammatory cells may switch to a proresolution M2 phenotype^51^. Henceforth, in disulfiram treated mice since infection is cleared, M2 polarizing factors like NOS2 (iNOS) were elevated than the doxycycline treated group and PBS infected group at day 21 or day 28 post infection (Figure 4). However, underlying mechanism involved in differential expression of chemo and cytokines and their effect on disease severity needs to be investigated.

Further, in this study we found lower bacterial burden in ear, heart and bladder of disulfiram treated mice compared to PBS treated infected mice at 21 days post infection (Figure 5), indicating that the disulfiram administration might have promoted the antibody mediated killing early in the infection, thus not only limit *B. burgdorferi* colonization in tissues but also altered the development of adaptive immune response, which may reduce the tissue inflammation as observed in heart^52,53^. In fact, *B. burgdorferi* infection leads to strong and sustained IgM response and delayed development of long-lived antibody and B cell memory^54^. So, disulfiram treated mice might have accelerated long lived antibody and B cell memory development, which resulted in statistically lower amount of total IgM, IgG and IgG1 in day 21 post infection (Figure 5). On the other hand later at day28 post infection, disulfiram treated mice have higher amounts of total IgM, IgG1 and IgG3isotypes, which all bind to C1q and activate the classical pathway, whereas IgG2a and IgG2b bind to the Fc receptor^55^. As such, it is likely possible that those immuno-complexes formed with C1q-binding antibodies cannot be opsonized by the complement system during infection due to the absence of C1q thus fail to be engulfed by phagocytes and accumulated within the circulation system. Lymphoadenopathy observed during Lyme borrreliosis is caused by a massive increase in lymph node cellularity triggered by the accumulation of live *B. burgdorferi* spirochetes into the lymph nodes. This increase in cellularity is due to accumulation of CD19+ B cells^33^. Disulfiram treatment not only alleviates lymphoadenopathy but also reduces the percentage of CD19+ B cells in day 28 post infected mice (Figure 6). An important function of CD4+ T cells is their ability to enhance antibody-mediated immunity by driving affinity maturation and the development of long-lived plasma cells and memory B cells. However, it appears unlikely that the protective B cell response to *B. burgdorferi*, a highly complex pathogen expressing many immunogenic surface antigens, is confined to T-independent antibody responses alone. Even though, disulfiram treated mice induces increase in percentage of CD3+ CD4+, Naïve, effector and memory T cells, further studies are needed to understand the role of these increased T cells in disease resolution and bacteria clearance.

In summary, the disulfiram drug not only successfully cleared the bacteria but also reduced the inflammation in heart tissue in C3H/HeN mice at day 28 post infection. Furthermore, disulfiram reduced antibody titers followed by nullifying lymphoadenopathy. The preclinical data offered here is beneficial in ascertaining the effectiveness of disulfiram and aids in future mechanistic and translation research studies. Moreover, the disulfiram can exploit multiple mechanisms to show its inhibitory effects both *in vitro* and *in vivo*. Although the results from our *in vivo* study cannot be extrapolated directly to clinical practice, they form strong basis for future follow-up studies, and promote the development of effective formulations of disulfiram for clinical management of Lyme disease.

## Materials and Methods

### Culturing and growth conditions of *B. burgdorferi* B31

*Borrelia burgdorferi sensu stricto* low passage strain B31 was (obtained from the American Type Culture Collection Manassas, VA) used for MIC tests and all infection studies in C3H/HeN mice. Bacteria cultures were started by thawing −80°C glycerol stocks of *B. burgdorferi* (titer, ∼10^7^ CFU/ml) and diluting 1:40 into fresh Barbour-Stoner-Kelly (BSK) complete medium with 6% rabbit serum followed by incubating at 33°C. After incubation for 4-5 days log phase, and 8-9 days stationary-phase *B. burgdorferi* culture (∼10^6^ borrelia/mL) was transferred into a 48-well plate for evaluation with the drugs.

### Drug formulations

The disulfiram (Sigma, USA) stock solution (50 mM) was made by dissolving in sterile 30% hydroxypropyl β-cyclodextrin (Sigma) and also another disulfiram stock solution (20 mM) was made by dissolving in sterile 100 % DMSO (Tocaris bioscience, UK). A stock solution of 100 mM of doxycycline (as a positive control) was made by dissolving the doxycycline powder in ultra-pure MilliQ water. All drug stocks were passed through 0.22 μm filters (Millipore-Sigma), used within 72 h of preparation and were not subject to freezing temperatures. Working solutions was made by mixing desired volume of stock solutions in desired volume of ultra-pure MilliQ water. Further, the vehicle for hydroxypropyl β-cyclodextrin (cyclodextrin) and DMSO controls were made similarly and it is important to note that the vehicle controls were identical to the test formulation in every single aspect except for the active ingredient. This measure was strictly followed for vehicle control wherever used in entire study.

### In-vitro testing of antibiotics by BacTiter Glo® assay, microdilution and SYBR Green I/PI assay methods

The MIC was determined by using Bac Titer-Glo microbial cell viability assay^29^. After 72 hours, 100 µL of culture was taken from each well and mixed with 100 µL of Bac Titer-Glo® reagent (Promega, Madison, WI, USA). Then, the assay was performed according to the manufacturer’s instructions. Luminescence was measured on a CLARIOstar micro plate reader at an integration time of 500 milliseconds.

A standard microdilution method was used to determine the minimum inhibitory concentration (MIC) of the antibiotics tested^56^. Approximately, 1 × 10^6^ *B. burgdorferi* were inoculated into each well of a 48-well tissue culture microplate containing 900 μL of BSK medium per well. The cultures were then treated with 100 μL of each drug at varying concentrations ranging from 0.625, 1.25, 2.5, 5, 10 and 20 μM. Control cultures were treated with respective vehicles, and all experiments were run in triplicate. The well plate was covered with parafilm and placed in the 33°C incubator with 5% CO_2_ for 4 days. Spirochetes proliferation was assessed using a bacterial counting chamber (Petroff-Hausser Counter) after the 4-5 days incubation followed by dark-field and fluorescence microscopy. To further determine the minimum bactericidal concentration (MBC) of the antibiotics tested (the minimum concentration beyond which no spirochetes can be sub cultured after a 3-week incubation period), wells of a 48-well plate were filled with 1 mL of BSK medium and 20 μL of antibiotic-treated spirochetes were added into each of the wells. The well plate was wrapped with parafilm and placed in the 33°C incubator with 5 % CO_2_ for 3 weeks (21 days). After the incubation period, the plate was removed and observed for motile spirochetes in the culture by dark-field and further cell proliferation was assessed using the SYBR Green I/PI assay fluorescence microscopy. All these experiments were repeated at least three times. Statistical analyses were performed using Student’s t-test.

### Dynamic light scattering

The stock solutions of 1M disulfiram was prepared either in DMSO or in 30% (w/v) hydroxypropyl ß-cyclodextrin (CD). Disulfiram was then diluted in bovine serum albumin (BSA) solution to obtain disulfiram concentration 0.125 µM, 0.25, 0.5, 10, 25, 50, and 100 µM, and 5% (w/v) BSA in the final solution for DLS. The measurements were obtained from Brookhaven 90-Plus particle size analyzer (Brookhaven instruments corporation) at an angle of 90° with 10% dust cutoff filter. The results represent average of three measurements.

### Atomic force microscopy

Atomic force microscopy (AFM) samples have been prepared from drugs disulfiram-CD and disulfiram-DMSO solutions of respective concentrations (100 μM, 25 μM, 10 μM and 5 μM) on clean silicon wafers that were plasma-treated to increase hydrophilicity. 10 µL droplets were deposited, spreading for most of the surface of 1 cm^2^ wafers and were quickly dried in a desiccator under vacuum to minimize additional aggregation due to local increase in concentrations. AFM imaging has been performed with NX-10 AFM (Park Systems, Korea) operating in non-contact mode with Micromasch NCS15 AL BS tips (NanoandMore, USA) at 0.8 Hz with 256 pixels per line.

### *In vivo* testing of drugs in immunocompetent C3H/HeN mice

Four weeks old female C3H/HeN mice, were purchased from Charles River Laboratories, Wilmington, Massachusetts. All mice were maintained in the pathogen-free animal facility according to animal safety protocol guidelines at Stanford University under the protocol ID APLAC-30105. All experiments were in accordance with protocols approved by the Institutional Animal Care and Use Committee of Stanford University. The mice (5 week) were infected subcutaneously close behind the neck with 0.1 mL BSK medium containing log phase 10^5^ *B. burgdorferi* B31. For *in vivo* studies, we have used only disulfiram soluble in cyclodextrin. On the 14 and 21 days post Bb infection, the mice were intraperitoneally administered a daily dose of drugs, disulfiram (75 mg/kg) and doxycycline (50 mg/kg) for 5 consecutive days (Figure 3A). After 48 hours of the last dose of administering compounds, both groups (day 21 and day 28 post Bb infection) of mice were terminated and their urinary bladders, ears, and hearts were collected. The DNA was extracted from urinary bladder, ear and heart. The absence of *B. burgdorferi* marked the effectiveness of the treatment in these organisms. Quantification of important pro/anti-inflammatory immune marker transcripts and histopathology of heart was also done. At termination on day 28 post infection, spleen and peripheral lymph nodes (axillary, brachial, cervical and inguinal) were also collected for immunophenotyping.

### Quantitative (Q-PCR) and Real-time PCR (RT-PCR) analysis

Urinary bladder, ear punches, heart bases were homogenized and DNA was extracted using the NucleoSpin tissue kit according to the manufacturer’s instructions (Düren, Germany). Q-PCR from above tissues were performed in blinded samples using *B. burgdorferi* Fla-B gene specific primers and a probe. These primers were listed as follows: Fla-B primers Flab1F 5’-GCAGCTAATGTTGCAAATCTTTTC-3’, Flab1R 5’-GCAGGTGCTGGCTGTTGA-3’ and TAMRA Probe 5’-AAACTGCTCAGGCTGCACCGGTTC-3’ according to the published protocol. Reactions were performed in duplicate for each sample. Results were plotted as the number of Fla B copies per microgram of tissue. The lower limit of detection was 10 to 100 copies of *B. burgdorferi* Fla-B DNA per mg of tissue. In addition to standard laboratory measures to prevent contamination, negative controls (containing PCR mix, Fla-B primers, probe, and Taq polymerase devoid of test DNA) were included.

Total RNA was extracted from tissues using RNeasy mini kit (Qiagen, USA) and reverse transcribed using a high-capacity cDNA reverse transcription kit (Invitrogen, USA). cDNA was subjected to real-time PCR using primer and TAMRA probes (Stanford Protein and Nucleic acid Facility) previously described^57^. PCR data are reported as the relative increase in mRNA transcript levels of CxCL1 (KC), CxCL2 (MIP-2), CCL5 (RANTES), IL-10, TNF-α, IFN-γ, iNOS and NOS2 normalized to respective levels of GAPDH.

### Histopathology

For histopathology, heart samples were fixed in 10% formalin buffer followed by staining of vertical histological sections with hematoxylin and eosin dye. Heart tissues were assessed for inflammation by microscopic examination at intermediate (10X), and high (40X)-power magnification and were scored for severity of inflammation (carditis, vasculitis) according to the percentage of inflammation at the heart base upon examination at low power (10X). Scores of 0 (none), 1 (minimal; less than 5%), 2 (mild; between 5% and 20%), 3 (moderate; between 20% and 35%), 4 (marked; between 35% and 50%), and 5 (severe; greater than 50%) were assigned for the severity of inflammation^58,31^. Myocarditis consisted of focal or diffuse interstitial infiltrates of mononuclear leukocytes in the myocardium. The microscopic photographs were captured on Olympus CX-41 microscope (Olympus, Tokyo, Japan). The images are shown at 10X and 40X magnification

### Quantification of total Immunoglobulins in serum by ELISA

Quantification of total mouse immunoglobulin concentration IgA, IgM, IgG, IgG1, IgG2a, IgG2b, and IgG3 in mouse serum was done using Ready-Set-Go ELISA kits (Invitrogen, USA).

### Flow cytometry

Single cell suspensions of lymphoid tissues were prepared as described^57^, live/dead cell viability stain was used to eliminate dead cells followed by single cells separation from doublets by FSC-A vs FSC-H plots. Cells were incubated in Fc blocking for 15 min at 4°C in staining buffer and incubated with the appropriate marker for surface staining in the dark for 30 min at 4°C. Surface lineage markers for T cells CD3, CD4, CD8, CD62L, CD44 and B cells was CD-19 conjugated with PercP Cy5.5 (Tonbo Biosciences) were described previously^57^. Cells were acquired on a BD-LSR II flow cytometer and data analyzed using Flow Jo software.

### Statistics

Data analysis was done using Graph Pad Prism software. Single comparisons within uninfected or drug treated groups and infected groups were analyzed with two-tailed paired t-test, with unpaired t-test with Welch’s correction and with multiple t-tests. α = 0.05 for all tests. *p < 0.05, ** p < 0.01, *** p < 0.001, ****p <0.0001.

## Acknowledgements

This work was accomplished with a generous support from the Bay Area Lyme Foundation. We thank Michal Caspi Tal and the Flow Cytometry Core in Institute for Stem cell biology and Regenerative medicine, Stanford University for providing access to FACS facility and also thank Mohammed Inayathullah from our BioADD lab for valuable suggestions regarding drug solubility and editing the manuscript.

## Author Contributions

HHSP performed the all experiments and analyzed the data. JB, a certified MD pathologist helped in histopathology studies, MI performed and analyzed DLS study, AVM performed AFM imaging, KMK helped in fluorescent imaging and *in vivo* studies. HHSP and JR designed the study and wrote the paper. All authors read and approved the final manuscript.

## Conflict of Interest

Jayakumar Rajadas are listed on the following patent titled “Methods and drug compositions for treating Lyme disease” under international patent application no WO2017124080A1. All other authors report no conflicts of interest in this work.

