## supplemental file 1 for "Repurposing disulfiram (Tetraethylthiuram Disulfide) as a potential drug candidate against *Borrelia burgdorferi in vitro and in vivo*"

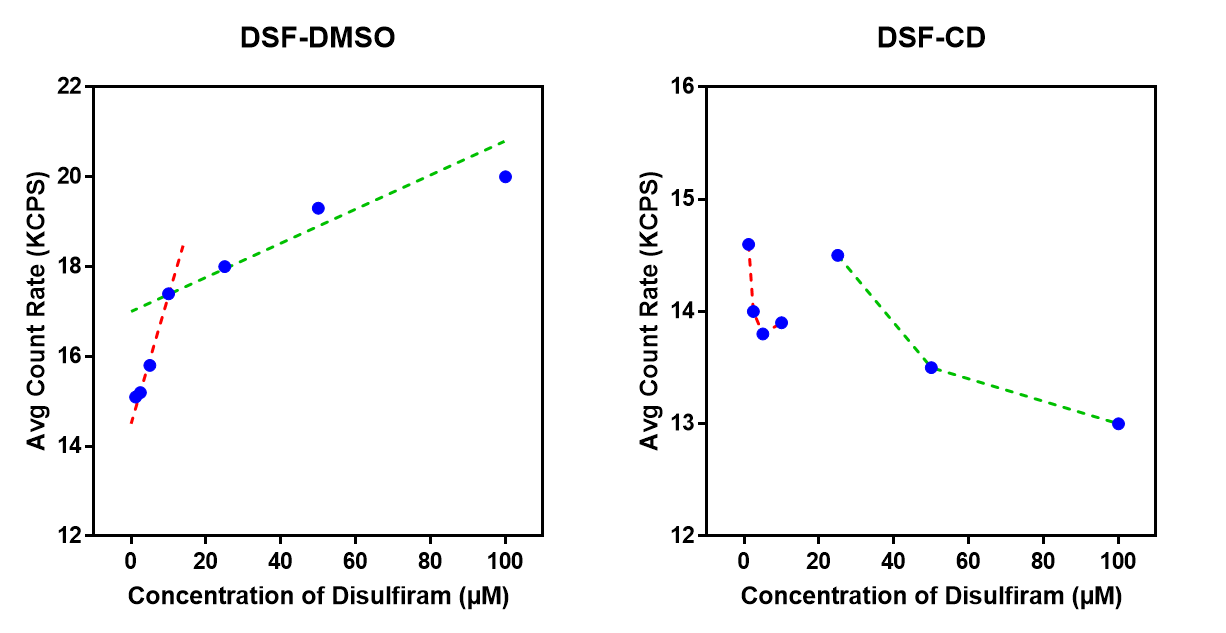

A

B

Supplemental figure 1 : Dynamic light scattering (DLS) analysis of disulfiram at different concentration in presence of 5% (v/v) bovine serum albumin (BSA) solution. The disulfiram solution was prepared by diluting the concentrated disulfiram stock solutions dissolved in either in **(A)** DMSO or in **(B)** hydroxypropyl ß-cyclodextrin (30% w/v) solution. The average count rate at different concentrations of disulfiram indicate two different aggregation patterns with a critical concentration at 10 µM.
