## supplemental file 2 for "Repurposing disulfiram (Tetraethylthiuram Disulfide) as a potential drug candidate against *Borrelia burgdorferi in vitro and in vivo*"

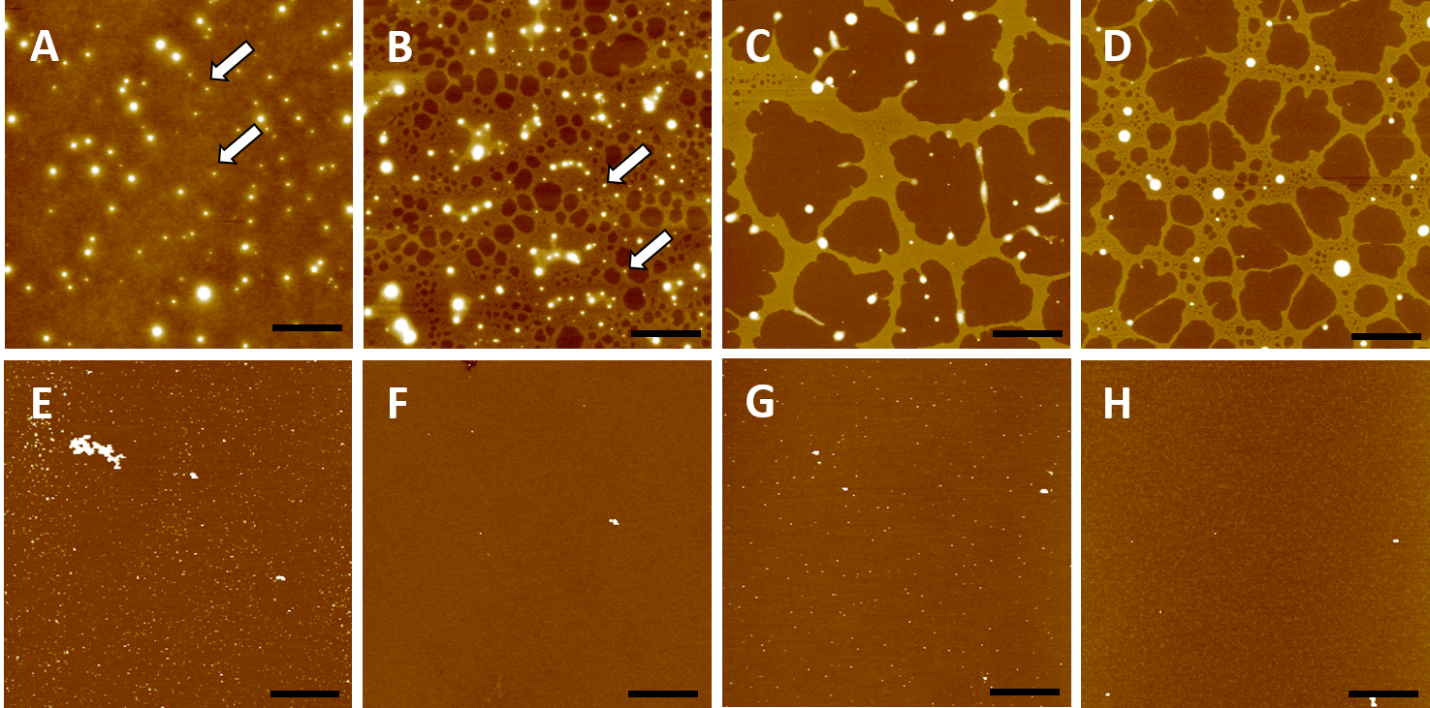

Supplemental figure 2: AFM images of disulfiram formulations prepared in cyclodextran **(A-D)** and DMSO **(E-H)** in the following solution concentrations: **A,E** – 100 µM, **B,F** – 25 µM, **C,G** – 10 µM, **D,H** – 5 µM. White arrows mark small size aggregates in the images which formed regardless of local aggregation due to solution drying during sample preparation. Vertical scale contrast is same for all images (4 nm). Scale bar is 2 µm.
