## supplemental file 3 for "Repurposing disulfiram (Tetraethylthiuram Disulfide) as a potential drug candidate against *Borrelia burgdorferi in vitro and in vivo*"

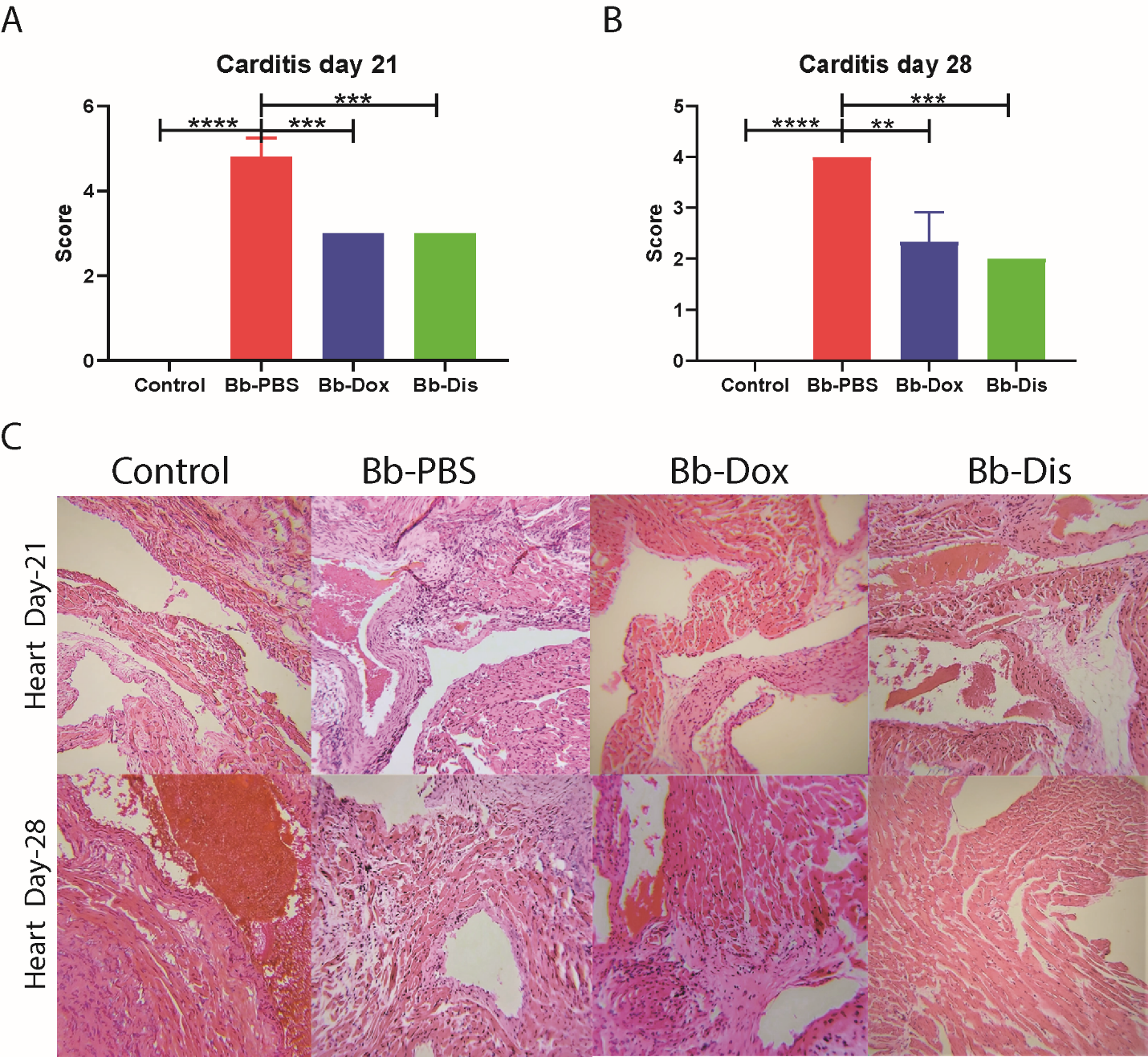

Supplemental figure 3: **A & B.** Histopathology was empirically quantified by scoring carditis blindly in 5 fields per sample and averaging per group. **C.** Photomicrographs (10X) of hematoxylin and eosin stained heart sections; arrows depict the mono nuclear leucocyte infiltrates. Statistics by unpaired t test with Welch’s correction between control versus infected and also between drug treated group versus infected group. ** p < 0.01, *** p < 0.001, ****p <0.0001.
